# A Novel Antenna Protein Complex in the Life Cycle of Cyanobacterial Photosystem II

**DOI:** 10.1101/660712

**Authors:** Daniel A. Weisz, Virginia M. Johnson, Dariusz M. Niedzwiedzki, Min Kyung Shinn, Haijun Liu, Clécio F. Klitzke, Michael L. Gross, Robert E. Blankenship, Timothy M. Lohman, Himadri B. Pakrasi

## Abstract

In oxygenic photosynthetic organisms, photosystem II (PSII) is a unique membrane protein complex that catalyzes light-driven oxidation of water. PSII undergoes frequent damage due to its demanding photochemistry. However, many facets of its repair and reassembly following photodamage remain unknown. We have discovered a novel PSII subcomplex that lacks five key PSII core reaction center polypeptides: D1, D2, PsbE, PsbF, and PsbI. This pigment-protein complex does contain the PSII core antenna proteins CP47 and CP43, as well as most of their associated low–molecular–mass subunits, and the assembly factor Psb27. Immunoblotting analysis, multiple mass spectrometry techniques, and ultrafast spectroscopic results supported the absence of a functional reaction center in this chlorophyll–protein complex. We therefore refer to it as the ‘no reaction center’ complex (NRC). Additionally, genetic deletion of PsbO on the PSII lumenal side resulted in an increased NRC population, indicative of a faulty PSII repair scheme at the cellular level. Analytical ultracentrifugation studies and clear native acrylamide gel analysis showed that the NRC complex is a stable pigment-protein complex and not a mixture of free CP47 and CP43 proteins. Our finding challenges the current model of the PSII repair cycle and implies an alternative PSII repair strategy. We propose that formation of this pigment-protein complex maximizes PSII repair economy by preserving an intact PSII core antenna shell in a single complex that is available for PSII reassembly, thus minimizing the risk of randomly diluting multiple recycling components in the thylakoid membrane following a photodamage event at the RC.

**Significance statement:** Photosystem II (PSII) converts sunlight into chemical energy, powering nearly all life on Earth. The efficiency of this process is maximized under various environmental conditions by a frequent repair and reassembly cycle that follows inevitable PSII damage even during normal oxygenic photosynthesis. We have isolated a novel pigment protein PSII subcomplex in which, surprisingly, the reaction center (RC) components of PSII are absent. Formation of this stable chlorophyll-protein complex suggests a protective mechanism whereby longer-lived PSII subunits are ‘unplugged’ from the damaged RC to prevent harmful, aberrant photochemistry during RC repair. This finding provides intriguing new insight into how PSII is assembled and rebuilt to optimize its performance to optimally catalyze one of the most challenging reactions in biology.

## Introduction

Photosystem II (PSII) is a large pigment-protein complex embedded in the thylakoid membranes of all oxygenic photosynthetic organisms: cyanobacteria, algae, and plants. PSII plays a central role in energy flow in the biosphere by harnessing sunlight to split water molecules into protons, electrons, and molecular oxygen, ultimately yielding the vital high-energy molecules ATP and NADPH. This process of converting solar energy to chemical energy powers nearly all life on Earth, while simultaneously producing the oxygen we breathe.

Crystal structures of functional PSII (1–6) have revealed that the water-splitting reaction is catalyzed by a Mn_4_CaO_5_ (Mn) cluster bound to the lumenal surface of PSII. D1 and D2, two ~30 kDa transmembrane proteins, form a heterodimer at the center of PSII. They coordinate the primary electron transport chain cofactors and contribute most of the Mn-cluster ligands. D1 and D2 associate with the smaller (<10 kDa) subunits PsbE and PsbF (α- and β-subunits of cytochrome *b*_559_) and PsbI. Together, these five proteins comprise the core “reaction center” (RC) complex, the smallest PSII subcomplex capable of light-induced charge separation (7). Surrounding the RC subunits are CP47 and CP43, two ~50 kDa proteins that bind chlorophyll *a* (Chl *a*) molecules and serve as antennae, harvesting and funneling light energy towards the RC to drive PSII photochemistry. Around 10 additional low-molecular mass (LMM) subunits bind to fully assembled PSII, contributing to the structural and functional optimization of the complex (8). Finally, functional PSII contains several membrane-extrinsic hydrophilic proteins (PsbO, PsbU, PsbV, and PsbQ in cyanobacteria) (9, 10), bound at the lumenal surface of the complex, that stabilize the Mn cluster.

PSII undergoes frequent oxidative damage owing to the demanding electron-transfer chemistry it performs (11–13). D1 is damaged and replaced most frequently of all proteins in the complex, closely followed by D2 (14–17). CP47 and CP43 are, however, more long lived (15, 17). This damage leads to partial disassembly of PSII, replacement of each damaged subunit with a new copy, and reassembly of PSII, in an intricate process known as the PSII repair cycle (14, 18, 19) (Fig. S1). This cycle operates concurrently with *de novo* PSII synthesis, which involves stepwise assembly of PSII from component subcomplexes. Many accessory proteins, such as Psb27 and Psb28 (20–24), bind exclusively to particular PSII subcomplexes to aid in a specific aspect of assembly. Much effort has been invested in characterizing the various subcomplexes that form during the PSII life cycle, but because they are typically found in low abundance and only form transiently, many details remain unclear.

In this study, we describe the identification of a novel PSII subcomplex from the cyanobacterium *Synechocystis* sp. PCC 6803 (*Synechocystis* 6803). This subcomplex specifically lacks the five RC subunits, and therefore, we refer to it as the ‘no-RC’ (NRC) complex. We discuss the implications of this complex in the context of the PSII life cycle and propose that its formation maximizes the efficiency of PSII repair and minimizes collateral damage to components of PSII that were unharmed following an initial photodamage event.

## RESULTS

### Isolation of a novel PSII subcomplex

We purified PSII complexes from the His47 and Δ*psb*O-His47 strains of *Synechocystis* 6803 by FPLC using a nickel affinity column. These complexes were then analyzed by high resolution clear native acrylamide gel electrophoresis (Fig. 1A). As expected, in the Δ*psb*O-His47 strain, a major band was seen that represents the PSII monomer (PSII-M), whereas no PSII dimer was found as it is not formed in this strain (10, 24). Additionally, a green band just below the monomer was observed that corresponds to the RC47 complex previously observed in the literature (14, 19). In the His47 strain, a major band was seen that represents the PSII dimer (PSII-D), while PSII-M and the RC47 bands were also present. Interestingly, an additional green band of unknown identity was present at a lower molecular weight than the monomer in each strain. In-gel digestion followed by tandem mass spectrometry (MS) indicated sharply decreased D1, D2, and PsbE content in the lower molecular weight band from both strains compared to PSII-M (Table S1), and presence of CP47 and CP43, though other assay methods were needed (see below) for more reliable quantitative information.

**Fig. 1.**
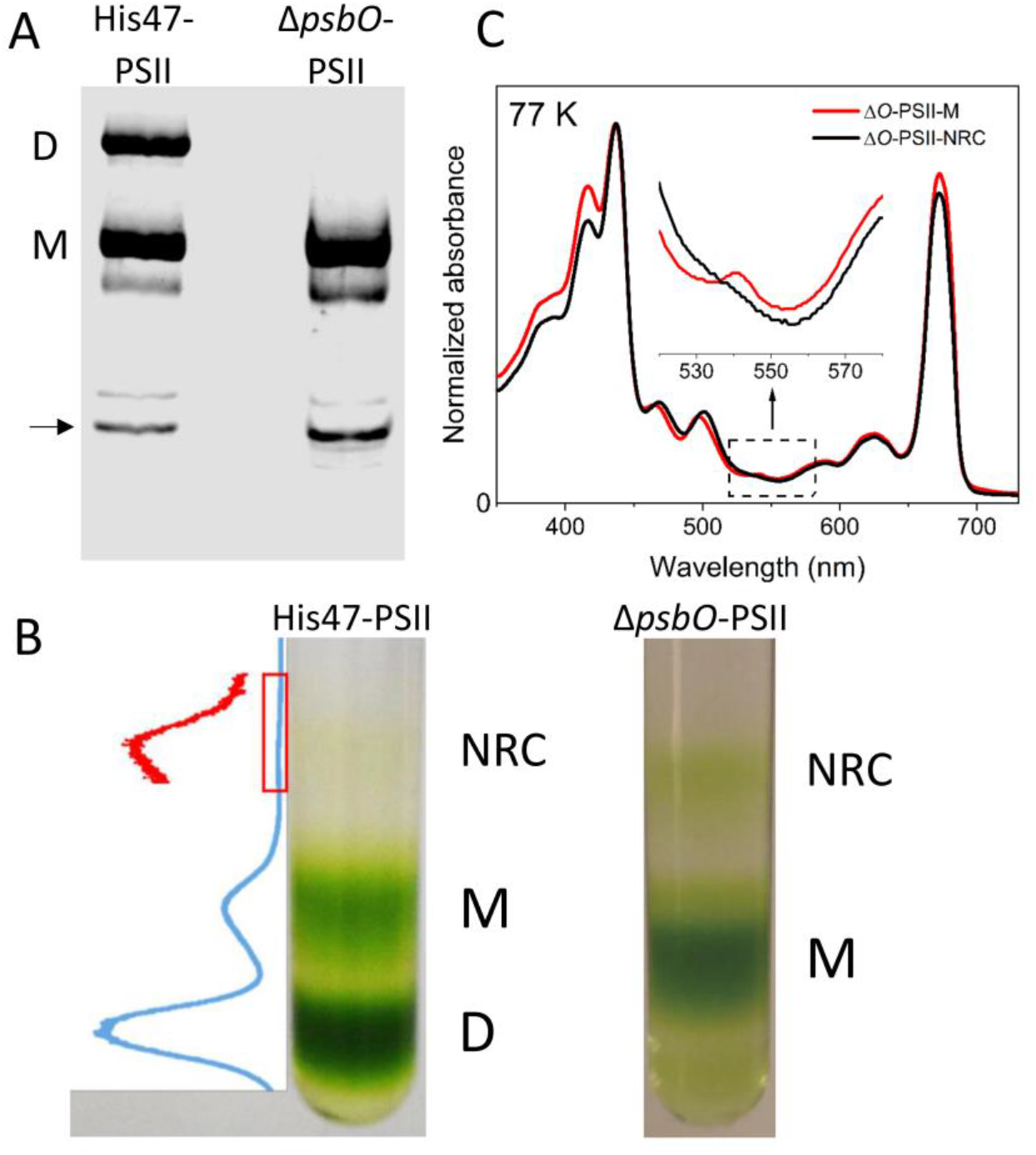
Isolation and characterization of a novel PSII subcomplex (NRC). (A) Clear Native PAGE of His-CP47-tagged PSII complexes from the His47 and Δ*psb*O-His47 strains. M, PSII monomer; D, PSII dimer. The arrow indicates the unknown band that was characterized further. (B) Green bands corresponding to the NRC complex and PSII monomer (M) formed following ultracentrifugation of His-CP47-tagged PSII samples in a 5-30% linear glycerol gradient from the His47 and Δ*psbO*-His47 strains. Pixel intensity (plotted as inverse of gray value computed in ImageJ (58) is shown on the left of the His47 gradient. The red curve shows the zoomed-in view of the region corresponding to the NRC band. (C) Steady-state absorption spectra of PSII-M and PSII-NRC at 77 K. To enable comparison, spectra were normalized to unity at the maximum of the Soret band (417 nm). The arrow shows a small peak visible in the spectrum of PSII-M that corresponds to the Q_x_ band of pheophytin *a* present in the PSII reaction center, and which is missing in the NRC spectrum.

To obtain a sufficient quantity for further characterization of the unknown complex, PSII preparations were then subjected to glycerol gradient ultracentrifugation to separate distinct His-CP47-containing complexes (Fig. 1B). Again as expected, in the Δ*psb*O-His47 strain, a major band was seen that represents the PSII monomer (PSII-M), whereas no PSII dimer was found as it is not formed in this strain (10, 24), and in the His47 strain, a major band was seen that represents the PSII dimer (PSII-D) in addition to the monomer band. Consistent with the results in the clear native gel, a lower molecular weight chlorophyll-containing band was observed in each strain. This band was harvested and concentrated. When the purified lower molecular weight band from the *ΔpsbO* strain was run next to *ΔpsbO*-PSII by clear native gel electrophoresis, it co-migrated with the band observed initially (Fig. S2). Tandem MS was performed after in-solution digestion of the complex from Δ*psbO* cells. The dominant PSII proteins were CP47 and CP43. According to the semi-quantitative information derived from such an experiment, levels of D1, D2, PsbE, and PsbF were markedly decreased compared to the corresponding PSII-M (Tables S2-S3), consistent with the results from in-gel digestion on the lower molecular weight band described above.

The 77-K absorption spectra of *ΔpsbO*-PSII-M and the novel complex were nearly identical (Fig. 1C), with the exception that a small peak around 540 nm corresponding to pheophytin *a* (25) was absent in the spectrum of the new complex. Pheophytin *a* is a cofactor in the PSII electron-transfer chain and is coordinated by D1 and D2. SDS-PAGE analysis showed no band detectable for the core RC proteins D1 and D2 (Fig. 2A). It, however, contained the inner antenna subunits CP47 and CP43, as well as Psb27, which binds to CP43 transiently during PSII assembly and repair (20, 21, 26, 27). In functional PSII, chlorophylls (Chl) bound to CP47 and CP43 harvest light and transfer excitation energy to Chl on D1 and D2, where primary PSII photochemistry occurs. A subcomplex containing both antenna proteins but lacking the two RC subunits to which they transfer energy was a surprising finding.

**Fig. 2.**
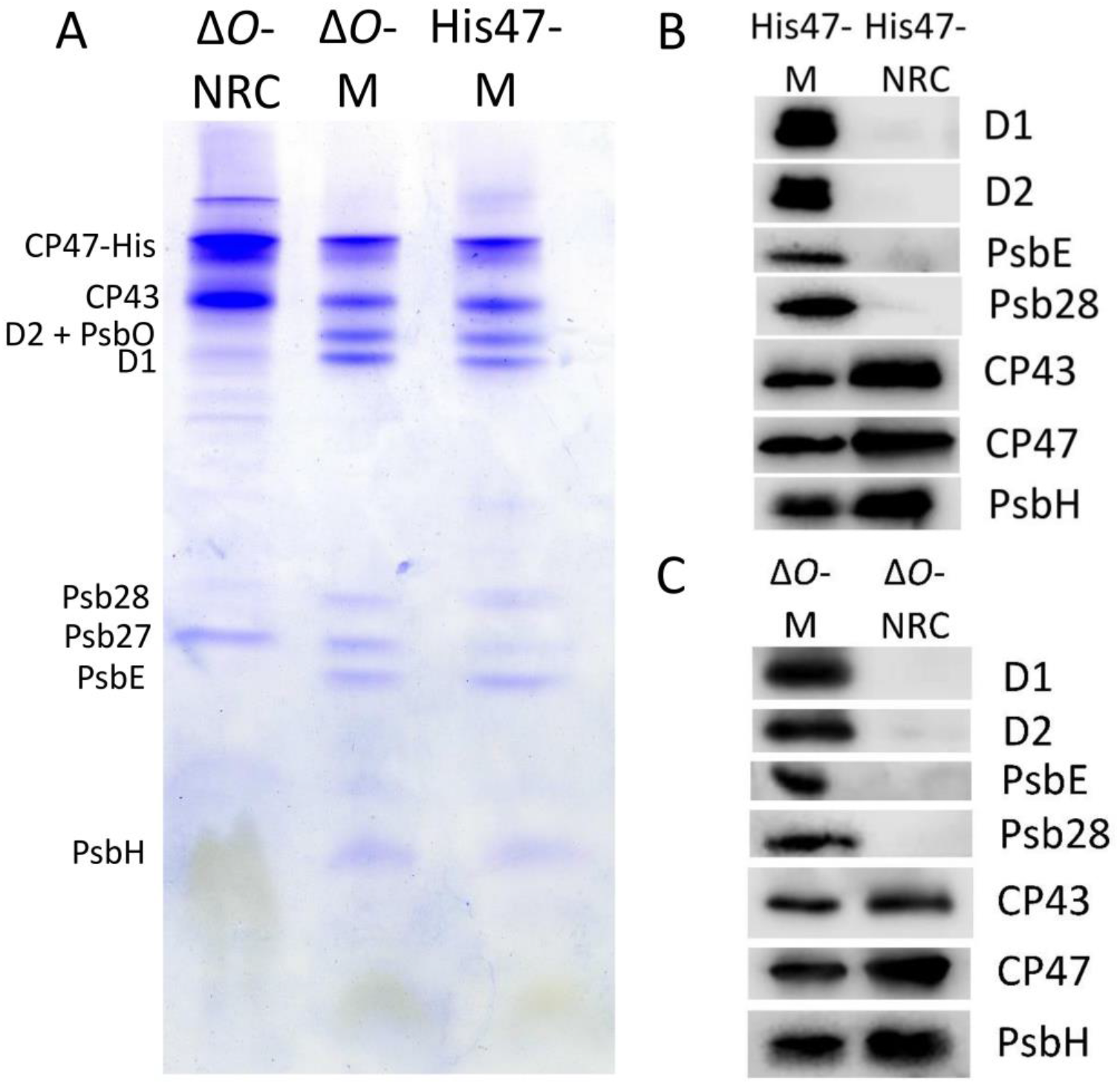
Major protein components of NRC. (A) SDS-PAGE analysis of PSII-M and PSII-NRC samples from the Δ*psbO*-His47 strain, and of PSII-M purified identically from the His47 strain. (B) Immunoblot analysis of PSII-M and PSII-NRC samples from the His47 strain. (C) Immunoblot analysis of PSII-M and PSII-NRC samples from the Δ*psbO*-His47 strain.

Consistent with SDS-PAGE, immunoblotting analysis (Figs. 2B, 2C) showed no detectable level of D1 and D2 in the novel complex isolated from both the *ΔpsbO* and His47 strains. Additionally, PsbE, another member of the PSII RC complex, was also absent (7). Presence of CP43 and CP47 in this complex, however, was confirmed, and the CP47-associated LMM subunit PsbH was detected as well. The MS results that showed severely decreased, but detectable, levels of D1 and D2 in the sample reflect the high sensitivity of the Q-Exactive mass spectrometry instrument and the inevitability of some impurities remaining in the sample following purification. The SDS-PAGE and immunoblot results show, however, that even if small amounts of D1 and D2 are present below their detection limits, they are not stoichiometric components of this complex.

For a more complete identification of the LMM subunits in this new complex, we employed MS to measure the mass of the intact protein components. We detected ten LMM subunits in the control *ΔpsbO*-PSII-M sample, whereas only seven of those ten were present in the new complex (Figs. 3, S3, Tables S4-S5). Remarkably, the three missing subunits are the three LMM components of the PSII RC: PsbE, PsbF, and PsbI. Taking our SDS-PAGE, immunoblot, and MS results together, we conclude that the novel subcomplex specifically lacks all five of the PSII RC components, D1, D2, PsbE, PsbF and PsbI, but contains the rest of the PSII subunits observed in the control PSII-M sample. We therefore refer to it as the “no-reaction center” (NRC) complex. The presence of NRC in the His47 strain, seen through native gel, glycerol gradient, mass spectrometry, and immunoblot characterization, demonstrates that this complex does not form as a result of the absence of PsbO in the Δ*psb*O-His47 strain. The experiments described below were, therefore, performed using NRC from Δ*psb*O-His47 cells owing to the higher yield obtainable in this strain.

**Fig. 3.**
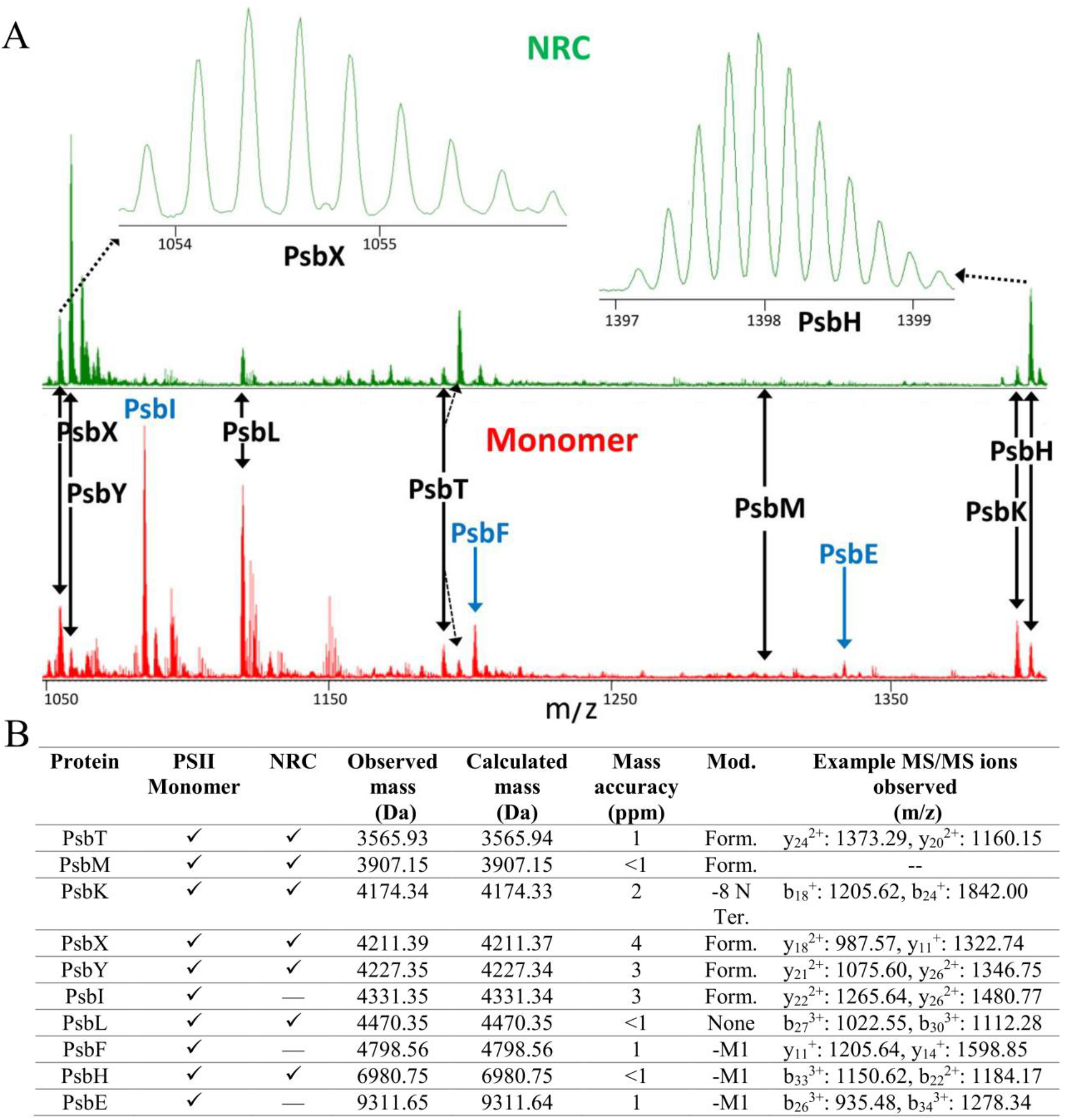
Mass spectra of the intact, low-molecular-mass subunits in Δ*O*-NRC (A, upper spectrum) and Δ*O*-PSII-M (A, lower spectrum) and summary table of the results (B). Most unlabeled peaks in the spectra correspond to additional charge states or oxidation products of labeled subunits. Subunits labeled in black afforded roughly comparable signal intensities in both samples; subunits labeled in blue (PsbE, PsbF, PsbI) were not observed in the NRC spectrum. Mod., modification; Form., formylation; −8 N-term., protein is cleaved after Alanine-8; -M1, loss of N-terminal methionine. See Fig. S3 for zoomed-in spectra of each subunit and comparison with theoretical spectra. MS/MS analysis was performed separately on each subunit (see Materials and Methods and Tables S4-S5) to confirm the identification, and example fragment ions are shown in (B). Mass accuracy of all MS/MS ions shown was ≤0.01 Da.

### Determination of the size of the NRC complex

Distributions of the Chl-containing species of the PSII-M and NRC samples were determined from sedimentation velocity experiments using analytical ultracentrifugation (AUC) (Fig. 4A). The NRC sample showed a major species at 6.1 S (peak b), the molecular weight (MW) of which was determined as 167.0 ± 5.3 kDa in a subsequent sedimentation equilibrium experiment (Fig. 4B). A minor species at 2.9 S was also observed (Fig. 4A), with MW 75.0 ± 3.5 kDa (Fig. 4B), which likely corresponds to dissociated CP47 (76 kDa) and/or CP43 (67 kDa) with their associated cofactors. For MW determination, the sedimentation equilibrium curves were analyzed using a two species fit based on the observation of two species from the sedimentation velocity experiments. The PSII-M sample showed a major species at 11.6 S (peak f), and three minor species at 2.9 S, 5.4 S, and 9.0 S. The MW of the major species was estimated as 436 kDa with a best-fit frictional coefficient ratio of 1.27, comparable to the MW of the PSII monomer determined in a previous study by Zouni and co-workers, using a similar method (28). The three minor species likely represent dissociation products.

**Fig. 4.**
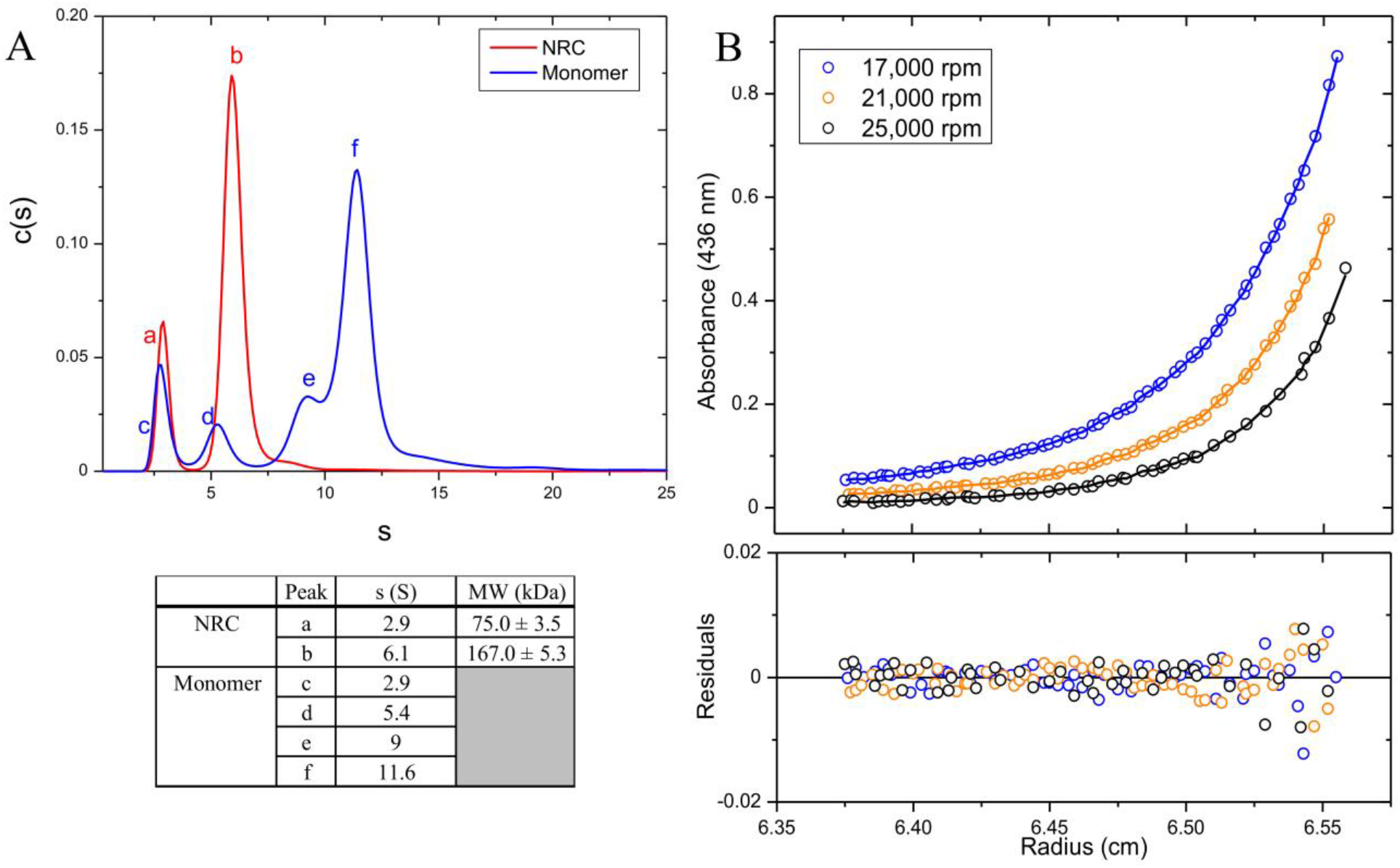
(A) Relative distribution of components in Δ*O*-PSII-NRC (red) and Δ*O*-PSII-M (blue) samples based on Svedberg coefficient (S), and (B) MW determination of the NRC complex from sedimentation equilibrium analysis. For experimental and calculation details, see “Materials and Methods.”

### The NRC complex lacks a functional reaction center

The absence of the RC proteins in NRC suggests that the complex cannot function as an RC. To test this hypothesis, we used ultrafast time-resolved fluorescence (TRF) measurements on the PSII-M and NRC samples. Both samples were excited at 625 nm, corresponding to a vibronic overtone of the Chl *a* Q_y_ band, and the Chl *a* fluorescence decay was monitored (Fig. 5A and B, shown as pseudo-color 2D profiles). The profiles revealed substantial differences between the two samples. The initial fluorescence profile of PSII-M, with a fluorescence maximum at ~680 nm, quickly evolved to a broader spectrum associated with another species with emission maximum at ~700 nm, consistent with energy transfer from antennae to a trap (the RC). Fluorescence spectra profiles of Chl *a* in NRC, however, remained relatively constant over time.

**Fig. 5.**
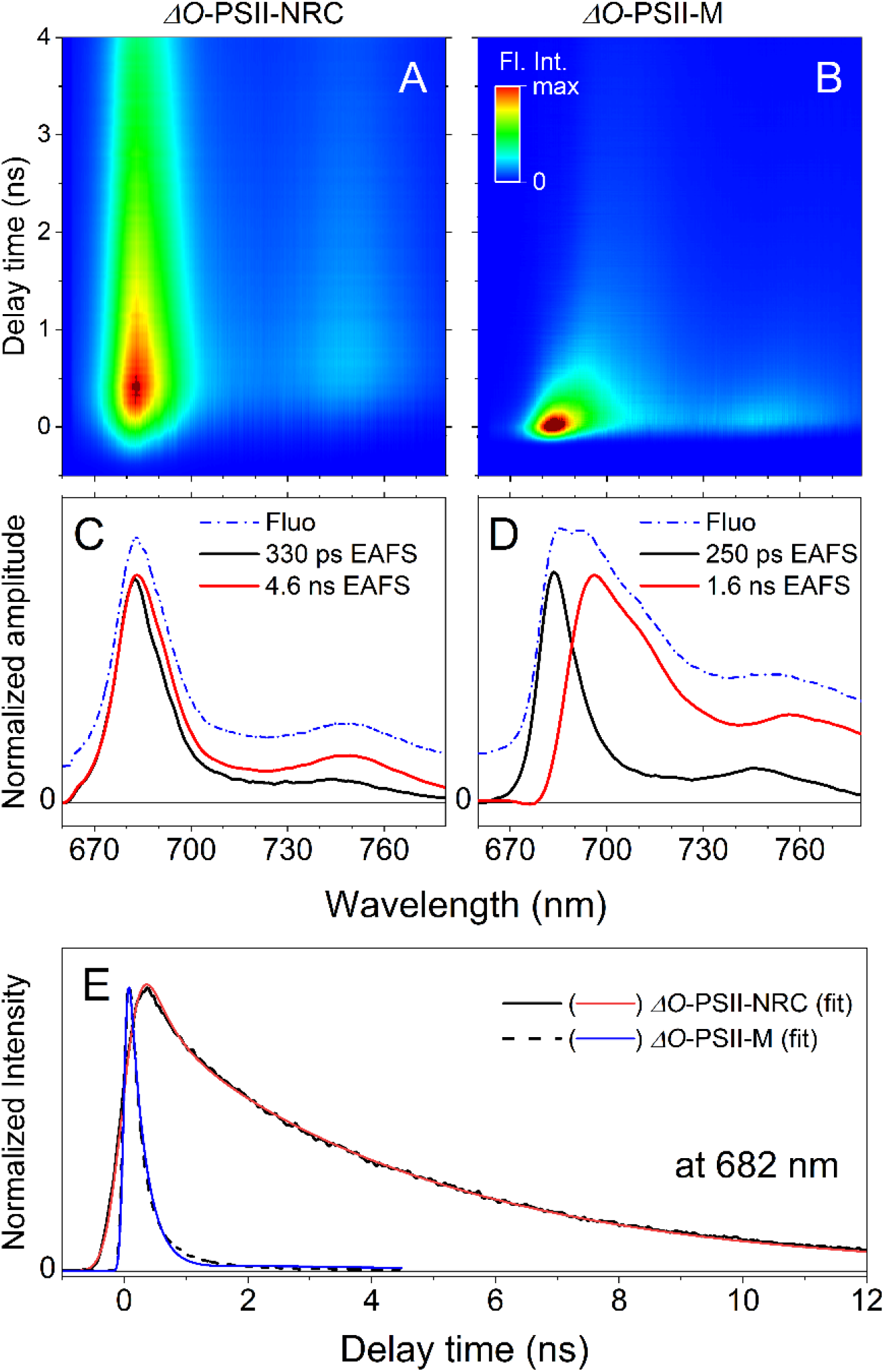
Time-resolved fluorescence of Δ*O*-PSII-NRC and Δ*O*-PSII-M samples. (A, B) 2D pseudo-color fluorescence decay profiles at 77 K upon excitation at 625 nm. (C, D) Global analysis results of TRF datasets. Dashed profiles (Fluo) correspond to time-integrated spectra and mimic steady-state fluorescence spectra. All profiles were normalized at their maxima for better comparability. (E) Example kinetic traces at 682 nm (maximum of fluorescence emission from CP47/CP43) accompanied with fitted traces from global analysis. Substantial shortening of fluorescence decay of the Δ*O*-PSII-M sample is associated with rapid excitation energy transfer from antenna into the PSII core. The different signal rise observed in the two traces originates from different temporal resolutions of time windows in which data were collected. EAFS – evolution associated fluorescence spectra.

The differences were further emphasized through kinetic analysis of the TRF data (Fig. 5C and D). Global analysis of the PSII-M and NRC spectra revealed two spectro-kinetic components (EAFS) for each sample (see Materials and Methods for more details). The faster PSII-M and NRC components, with lifetimes of 250 and 330 ps, respectively, resemble each other and correspond to excitation equilibration within the protein. The slower PSII-M component, with a lifetime of 1.6 ns, was red-shifted and reflects excitation-energy transfer from CP47 and CP43 to deep energetic traps in the RC. These PSII-M results closely mimic those obtained for PSII core complexes from the cyanobacterium *Thermosynechococcus vulcanus* under similar conditions (29). The slower NRC component, however, with a lifetime of 4.6 ns, was very similar to the fast NRC component, suggesting that both originate from the same species. The lifetime of 4.6 ns closely resembled that obtained for isolated CP47 and CP43 proteins (30), implying that no further excitation energy transfer occurs in the NRC complex, and this component corresponds to intrinsic decay of excited Chl *a*. Overall, these results demonstrated that excitation energy in PSII-M that was initially localized on Chl *a* bound to CP47 and CP43, was efficiently transferred to the RC. In contrast, no such RC energy trap exists in the NRC complex, and excitation energy remained localized on CP47 and CP43 until it decayed intrinsically.

## DISCUSSION

In this study, we discovered and characterized the “no-reaction center” (NRC) complex, a PSII subcomplex that contains the inner antenna subunits CP43 and CP47, Psb27, and seven LMM subunits, but is missing the five subunits that comprise the PSII RC: D1, D2, PsbE, PsbF, and PsbI. An assembled complex without all five RC components is remarkable and has not been described before. Through analytical sedimentation experiments, we determined the MW of the NRC complex to be 167.0 ± 5.3 kDa. This value is in between the sum of the apo (153 kDa)- and holo (194 kDa)-masses of the NRC proteins, indicating that some NRC cofactors with D1 or D2 interfaces may be destabilized in their absence. The determined MW is also close to the sum of the apparent masses of the individual CP43 and CP47 pre-complexes (around 180 kDa) measured previously (31). These results are not consistent with the NRC sample comprising a mixture of separate CP43 and CP47 pre-complexes, as each of these pre-complexes have MWs roughly half of the determined MW. We can also rule out homodimers of the CP43 and CP47 pre-complexes. Given the respective sizes of the two pre-complexes (31), such dimers would differ in mass by around 40 kDa, not consistent with the observed single, narrow, symmetrical major peak (Fig. 4, peak “b”) indicating a single species. Homodimers were also not detected during previous characterization of the pre-complexes (31). Overall, the analytical sedimentation results indicate that the NRC complex is a single species likely composed of a single copy of each of the PSII subunits identified in it.

The LMM subunits identified in NRC are consistent with the PSII crystal structures (2, 4, 5) in that all share interfaces with either CP43 or CP47 (except PsbY, which interfaces with another NRC subunit, PsbX). PsbH, PsbL, and PsbT were previously identified in the CP47 pre-complex, and PsbK in the CP43 pre-complex (31), consistent with our results. The presence of Psb27 is reasonable because it binds to CP43 (20, 27, 32), and the absence of Psb28 (Figs. 2B, 2C) is also reasonable as Psb28 associates closely with PsbE and PsbF (24) and would not be expected to bind in their absence.

In the crystal structures of mature PSII, D1 and D2 bridge the gap in the transmembrane region between CP43 and CP47. In their absence, the NRC subunits do not form a continuous structure, except for a very small overlap between PsbL and PsbT at the cytosolic surface. A significant CP43-CP47 interface would likely be necessary to preserve the structural integrity of this complex and could occur with CP43 and CP47 approaching each other and at least partially closing the gap left by D1 and D2.

Within the PSII life cycle, NRC could form during *de novo* synthesis and/or during the repair cycle (Fig. 6). During *de novo* synthesis, the complex would form as a spontaneous merger between newly synthesized CP43 and CP47 pre-complexes, from which CP47 followed by CP43 would attach to nascent RC complexes according to the current model (14, 33). The low abundance of NRC in the wild-type strain has likely prevented its detection until now. Many studies have used mutants that accumulate early assembly intermediates, but these valuable studies have often used either a ΔCP43 or a ΔCP47 strain (27, 34, 35), further preventing detection of the complex.

**Fig. 6.**
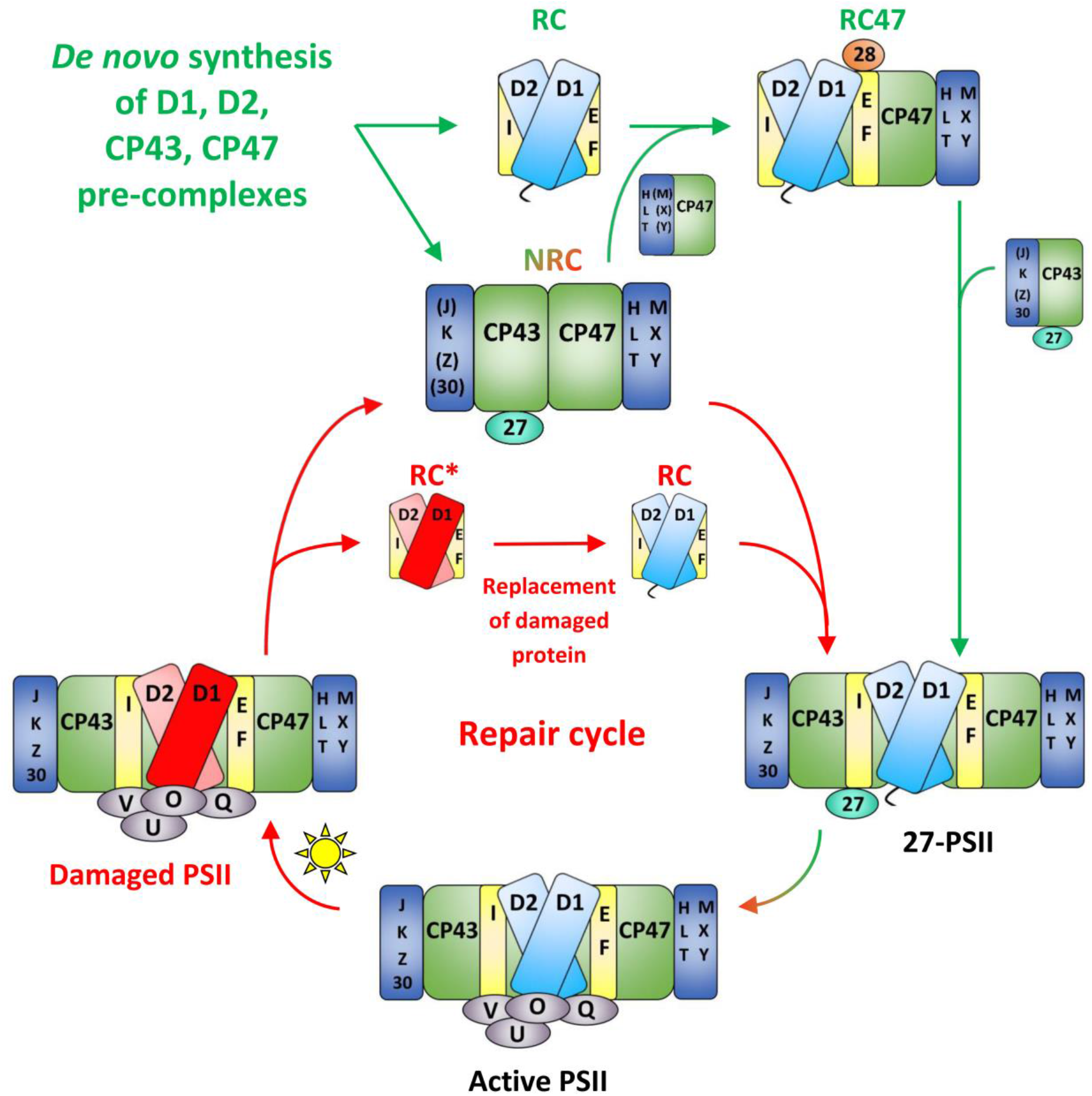
Schematic of the PSII life cycle based on previous findings (reviewed in (14, 18, 33, 36) and current results, focusing on the proposed positioning of the NRC complex. NRC may form during *de novo* synthesis (upper green arrows), during the repair cycle (red arrows), or both. Red D1, damaged version of D1; light red D2, slightly less frequently damaged version of D2. In a given RC* complex, either D1, D2, or both may be damaged. Several intermediary stages in the life cycle are omitted for clarity; e.g., after damage, PsbO, PsbU, PsbV, and PsbQ dissociate (36), Psb27 rebinds (21), and the complex monomerizes (36) before the proposed NRC formation step. Black text and red-green coloring represent steps common to both *de novo* synthesis and repair.

The second possibility, not mutually exclusive with the first, is that NRC forms during the repair cycle (Fig. 6), after PSII undergoes photodamage (most often at the D1 subunit). The repair steps are: 1) dissociation of the extrinsic proteins PsbO, PsbU, PsbV, and PsbQ (36), 2) rebinding of Psb27 (21), 3) monomerization of the complex (36), 4) an undefined further dissociation step (19), 5) D1 degradation (37–39), 6) insertion of a new D1 copy (14), and 7) PSII reassembly (14).

Within this known framework, NRC formation would most logically occupy step 4, the undefined dissociation step, with the damaged Psb27-PSII monomer splitting into RC and NRC components (Fig. 6). An implication is that D1 replacement would occur at the level of the RC complex (Fig. 6), not at RC47 as has been previously proposed (Fig. S1) (14, 19, 40). Meanwhile, CP43 and CP47 would be sequestered efficiently in one location, ready to be recycled back into PSII after repair. Though D1 is turned over most frequently, D2 has the second-highest turnover rate, approximately double that of CP43 and over three times that of CP47 (15). Dissociation of PSII into RC and NRC complexes following photodamage would allow maximal access for proteases to both D1 and D2, whereas, by interacting closely, CP43 and CP47 would shield each other from such protease attack (41). The recent interesting finding that PSII can remain stable and fully intact following Mn cluster removal (42) supports the notion that there may be an important efficiency benefit to maintaining a PSII subcomplex, once assembled, in as intact a state as possible, while still allowing for necessary repair to occur.

Damaged PSII is a liability because aberrant PSII photochemistry could result in further oxidative damage to the complex. During PSII assembly, D1 processing is a checkpoint that prevents premature formation of the highly oxidizing Mn cluster, which could result in damage to nearby subunits (26). The dissociation of the extrinsic proteins and the rebinding of Psb27 following photodamage likely achieves the same goal: inactivation of the Mn cluster at a time when it could only cause further damage. Perhaps the formation of the NRC complex can be understood in a similar manner. In addition to being an efficient disassembly/reassembly mechanism, separating into RC and NRC complexes following initial damage to D1 or D2 could serve a twofold protective purpose: 1) it lowers the total excitation energy reaching the RC by “unplugging” it from the CP47 and CP43 antenna subunits, and 2) it prevents aberrant PSII photochemistry from causing collateral damage to the NRC components, which are unlikely to have been harmed in the initial photodamage event. Future structural studies of the NRC Chl-protein complex are expected to shed light on the dynamic process during the PSII repair cycle.

## Materials and Methods

### Cell culture, PSII purification and protein analysis

Generation of the Δ*psb*O-His47 strain was reported previously (10). The HT3 (His47) strain was a kind gift from Dr. Terry Bricker (Louisiana State University, Baton Rouge, LA) (43). Cyanobacterial strains were grown in BG11 medium at 30 °C under 40 μmol photons m^−2^ s^−1^. The growth media were supplemented with 10 μg/mL spectinomycin and 5 μg/mL gentamicin (Δ*psb*O-His47 strain) or 5 μg/mL gentamicin (His47 strain). Histidine-tagged PSII complexes were purified by FPLC, as described previously (44), with minor modifications, and were stored in 25% glycerol (wt/vol), 10 mM MgCl_2_, 5 mM CaCl_2_, 50 mM MES buffer pH 6.0.

Following FPLC purification, PSII complexes were purified further by glycerol gradient ultracentrifugation, performed as described previously (24). To prepare one centrifuge tube, 0.6 mL 50% glycerol in RB buffer pH 6.5 was layered onto the bottom of the tube. The gradient was made from 6 mL each of stock solutions of 5% and 30% glycerol in RB buffer pH 6.5 containing 0.04% β-dodecyl maltoside. The stock solutions were added into the two chambers of a gradient maker connected to a peristaltic pump and were allowed to mix gradually as the pump delivered the mixture into the tube just below the top of the 50% glycerol layer. PSII sample containing 100 μg Chl *a* was diluted 1:5 into RB buffer pH 6.5 (final concentration 5% glycerol), then loaded onto the top of the gradient. Ultracentrifugation was performed at 180,000 *g* overnight at 4 °C. After centrifugation, the bands were harvested and concentrated by using Vivaspin 500 centrifugal concentrators (50 kDa cutoff) (Vivaproducts, Littleton, MA). The NRC complex was obtained in this manner consistently from numerous distinct biological preparations. Protein electrophoresis was performed as described previously (45, 46). For immunoblotting, gels were transferred onto a PVDF membrane (MilliporeSigma, Burlington, MA) followed by probing with specific antisera. Immunoblot imaging was performed with chemiluminescence reagents (MilliporeSigma, Burlington, MA) on a LI-COR Odyssey Fc (LI-COR Biotechnology, Lincoln, NE).

### Clear native polyacrylamide gel electrophoresis

High resolution clear native PAGE was performed as described in (47). 1.5×160×160 mm clear native polyacrylamide gel, 4-13%, was prepared using a gradient maker. Protein samples containing 10 μg of Chl *a* were loaded in each lane. Gel was run at 4mA for 16 hours at 4° C. Gels were imaged using a Li-COR Odyssey Fc (LI-COR Biotechnology, Lincoln, NE) using the 700 nm channel to visualize chlorophyll fluorescence.

### In-gel digestion and LC-MS/MS analysis

Protein samples were excised from the clear native polyacrylamide gel, destained and dehydrated with (1:1) 100mM ammonium bicarbonate:acetonitrile and digested in-gel with 13 ng/uL trypsin (Sigma) in 1mM TEABC. Following digestion, sample was extracted in 1% formic acid and subjected to LC-MS/MS.

Aliquots (5 μL, ~100 pmoles) of the peptide samples were separated online using a Dionex Ulimate 3000 RSLCnano pump and autosampler (Thermo Fisher Scientific, Waltham, MA, USA) and a column packed in-house utilizing ProntoSIL C18AQ, 3 μm particle size, 120 Å pore size (Bischoff, Stuttgart, Germany), in a 75 μm X 15 cm capillary. The mobile phase consisted of A: 0.1% formic acid in water and B: 0.1% formic acid in 80% acetonitrile/20% water (Thermo Fisher Scientific, Waltham, MA, USA). At a flow rate of 500 nL/min, the gradient was held for 5 min at 2% B and slowly ramped to 17% B over the next 30 min, increasing to 47% B over the next 30 min and then finally increasing to 90% B over 30 min and held at 90% for 10 min. The column was then allowed to re-equilibrate for 60 min with a flow of 2% B in preparation for the next injection.

The separated peptides were analyzed on-line by using a Q-Exactive Plus mass spectrometer (Thermo Fisher Scientific, Waltham, MA, USA) operated in standard data-dependent acquisition mode controlled by Xcalibur version 4.0.27.19. Precursor-ion activation was set with an isolation width of *m*/*z* 1.0 and with two collision energies toggled between 25 and 30%. The mass resolving power was 70 K for precursor ions and 17.5 K for product ions (MS2).

The raw data were analyzed using PEAKS Studio X (version 10.0, Bioinformatics Solution Inc., Waterloo, ON, Canada, www.bioinfor.com) and Protein Metrics Byonic and Byologic (Protein Metrics Inc., Cupertino, CA, www.proteinmetrics.com) (48). PEAKS was used in the *de novo* mode followed by DB, PTM, and SPIDER modes. Search parameters included a precursor-ion mass tolerance of 10.0 ppm and a fragment-ion mass tolerance of 0.02 Da. Variable modifications included all built-in PTMs. The maximum allowed modifications per peptide were 3; and the maximum missed cleavages were 2; false discovery rate, 0.1%. SPIDER (function) was used to identify unknown spectra by considering homology searches, sequence errors, and residue substitutions to yield a more confident identification.

Byonic searches employed the same database but used a precursor ion mass tolerance of 20 ppm and a fragment ion mass tolerance of 60 ppm with a maximum of 2 missed cleavages. Wildcard searches of ± 200 Da were employed to look for modifications in addition to regular PTM analysis. Protein false discovery rate threshold was determined by the score of the highest ranked decoy protein identified. All of the search results were combined in Byologic for validation.

### In-solution digestion and LC-MS/MS analysis

Samples each containing 2 μg Chl *a* of Δ*psbO*-PSII-M and Δ*psbO*-NRC were precipitated using the 2D Cleanup Kit (GE Healthcare, Chicago, IL) according to the manufacturer’s instructions. The pellets were resuspended in 20 μL 8M urea, 50 mM ammonium bicarbonate (ABC). Lys C was added at a 1:50 w/w protease:sample ratio and incubated at 37 °C for 2 hours. Samples were diluted to 1M urea with 50 mM ABC and trypsin was added at a 1:50 w/w protease:sample ratio. The samples were incubated overnight at 37 °C, then acidified to 1% formic acid and centrifuged to remove any insoluble material.

Aliquots (5 μL) of the digests were analyzed by LC-MS/MS as described in (24) with the following modifications: the LC gradient ran from 2-90% Solvent B with the following steps-12 min at 2% B, 33 min linear increase to 15% B, 30 min linear increase to 50% B, 15 min linear increase to 90% B, 9 min hold at 90% B, 1 min linear decrease to 2% B, and 30 min hold at 2% B; scan range was m/z 380-2200; for data-dependent MS/MS scans, resolution was 35,000 for ions at m/z 200 and AGC target was set at 2 × 10^5^ ions; and the top 10 ions were fragmented by HCD.

The raw data were loaded into PEAKS (version 8.5, Bioinformatics Solution Inc., Waterloo, ON) for protein identification. The data were searched against a database of the *Synechocystis* 6803 proteome using the built-in fusion decoy database for false discovery rate calculation (49). Search parameters were as follows: precursor-ion mass tolerance, 10.0 ppm; fragment-ion mass tolerance, 0.02 Da; variable modifications, all built-in modifications; maximum variable modifications per peptide, 3; maximum missed cleavages, 2; maximum nonspecific cleavages, 0; false discovery rate, 0.1%.

### Mass spectrometry of intact proteins

Δ*O*-M and Δ*O*-NRC samples, each containing 1.4 μg Chl *a*, were precipitated using the 2D Cleanup kit (GE Healthcare, Chicago, IL), following the manufacturer’s instructions. The pellets were resuspended in 100 μL 70% acetone, 19% water, 10% isopropanol, 1% formic acid v/v (31), diluted 1:3 in the resuspension solution, and infused directly into a Synapt G2 HDMS mass spectrometer (Waters, Milford, MA) at a flow rate of 500 nL/min by means of a syringe pump PHD Ultra (Harvard Apparatus, Holliston, MA). The mass spectrometer, equipped with a nanoelectrospray ionization source, was operated in sensitive ‘V’ mode with 10,000 mass resolving power (full-width at half-maximum). Positive-ion formation was achieved by applying a capillary voltage of 2.6 kV. The sampling and extraction cone voltages were 40 and 2 V, respectively, and the source temperature was 30 °C. Mass spectra were acquired between *m*/*z* 100 and 4,000 with an acquisition rate of one spectrum per sec. Data processing was with MassLynx 4.1 (Waters, Milford, MA). Final spectra were the average of 300 spectra collected in profile mode and converted to centroid data.

For LC-MS/MS analysis, ΔO-M and ΔO-NRC samples were prepared as described above for direct infusion. An aliquot (5 μL, ~1-2 pmol of intact proteins) was loaded onto a trap column (180 μm × 2 cm, C18 Symmetry, 5 μm, 100Å, Waters, Milford, MA) using solvent A (water with 0.1% formic acid). Peptides were eluted from a reverse phase C18 column (ProntonSIL 3μm, 120Å) 75 μm × 150 mm by increasing the fraction of solvent B (80% acetone, 20% water, and 0.1% formic acid). The gradient was supplied by a Dionex Ultimate 3000 instrument (Thermo Scientific, Inc., Sunnyvale, CA) and run from 2 to 17% solvent B over 65 min and then to 40% solvent B for 15 min, and then to 90% solvent B for 15 min at a rate of 500 nL/min followed by 15 min re-equilibration step with 98% solvent A. The Q-Exactive Plus spectrometer (Thermo Fisher Scientific, Waltham, MA, USA) was operated in standard mode with an inclusion list of all the LMM subunits observed by direct infusion on the Synapt G2 (Table S4). Peptide mass spectra (*m*/*z* range of 350-2000) were acquired at a high mass resolving power (7000 for ions at *m*/*z* 200) with the Fourier transform (FT) mass spectrometer. Precursor activation in HCD was performed with an isolation width of *m*/*z* 1.0 and a normalized collision energy of 30%. Default charge state was 3+ with charge exclusion of 1 and >8.

The LC-MS/MS raw data were submitted to the Protein Metrics (PMI) software package for analysis (48). Database searching was performed by the Byonic software using the *Synechocystis* 6803 phycobilisome and reaction centers protein database and a decoy database containing reversed protein sequences. Search parameters were: precursor ion mass tolerance 20 ppm, fragment ion mass tolerance 60 ppm, 0 cleavage sites, common post-translational modifications, and automatic peptide score cut. Protein false discovery rate threshold was determined by the score of the highest ranked decoy protein identified. The search results were combined in the Byologic software for validation and extraction of ion chromatograms with a mass window of 20 ppm. Manual quality control was performed using XCalibur Qual Browser assisted by the Protein Prospector package (http://prospector.ucsf.edu/prospector/mshome.htm) and Magtran (50).

### Ultrafast time-resolved fluorescence spectroscopy

All spectroscopic measurements were carried out at 77 K, using a VNF-100 liquid nitrogen cryostat (Janis, USA). The samples (Δ*O*-M and Δ*O*-NRC) were diluted in 60:40 v/v glycerol:RB buffer with 0.04% DM, that after cooling, formed fully transparent glass. Steady-state absorption spectra were recorded on a UV-1800 spectrophotometer (Shimadzu). Time-resolved fluorescence (TRF) imaging was performed using a universal streak camera system (Hamamatsu Corporation, Japan) based on N51716-04 streak tube and A6365-01 spectrograph from Bruker Corporation (Billerica, MA) coupled to an ultrafast laser system, described previously (51). The repetition rate of the exciting laser was 4 MHz, corresponding to ~250 ns between subsequent pulses. The excitation beam was depolarized, focused on the sample in a circular spot of ~1 mm diameter and set to a wavelength of 625 nm and very low photon flux of ~10^10^ photons/cm^2^ per pulse. The emission was measured at a right angle to the excitation beam with a long-pass 645-nm filter placed at the entrance slit of the spectrograph. The integrity of the samples was examined by observing the photon counts in real-time over the time course of the experiment. These were constant, indicating no detectable sample photodegradation. Prior to further analysis, all TRF datasets were subjected to singular value decomposition (SVD), a least-squares estimator of the original data leading to significant noise reduction (52).

The data were globally fitted with application of a simple fitting model that assumes irreversible direction of excitation decay from fastest to slowest decaying states. It is commonly called a sequential model, and spectro-kinetic components obtained from this fitting are typically called evolution-associated spectra (EAS). This nomenclature was adopted for the TRF analysis and the fitting results of those data are called evolution associated fluorescence spectra (EAFS) (53).

### Analytical sedimentation

After sample concentration following glycerol gradient ultracentrifugation (see above), Δ*O*-M and Δ*O*-NRC samples were buffer exchanged into RB buffer pH 6.5 containing 5% glycerol. Sedimentation velocity experiments were performed with an Optima XL-A analytical ultracentrifuge and An50Ti rotor (Beckman Instruments, Fullerton, CA) at 42,000 rpm (25 °C) as described previously (54). The experiment was performed at 4 and 13 μg/mL Chl *a*, while monitoring absorbance at 437 nm, with the results consistent at both concentrations. Data were analyzed using SEDFIT (55), to obtain c(s) distributions. The c(s) distribution function defines the populations of species with different sedimentation rates (sizes) and represents a variant of the distribution of Lamm equation solutions (55). The density and viscosity of the RB buffer at 25° C were determined using SEDNTERP. 0.76 mL/g was used as the partial specific volume (28).

Sedimentation equilibrium experiments were performed with NRC sample at the three indicated speeds (25 °C) starting at the lowest and finishing at the highest speed as described previously (56). 110 μL of the NRC sample at the same concentration used for the sedimentation velocity experiments and 120 μL of the buffer were loaded to an Epon charcoal-filled six-channel centerpiece. Absorbance data were collected at intervals of 0.003 cm in the step mode with five averages per step. Data were edited using SEDFIT to extract concentration profiles from each chamber and analyzed using SEDPHAT with the Species Analysis with Mass Conservation model (57).

## Acknowledgements

We thank Dr. Alexander Kozlov for helpful discussions. This work was supported by the Chemical Sciences, Geosciences, and Biosciences Division, Office of Basic Energy Sciences, Office of Science, U.S. Department of Energy (DOE) (Grant DE-FG02-99ER20350 to H.B.P.); the Photosynthetic Antenna Research Center, an Energy Frontier Research Center funded by the U.S. DOE, Office of Basic Energy Sciences (Grant DE-SC 0001035 to H.B.P., R.E.B., and M.L.G.); NIH Grant P41GM103422 to M.L.G. and NIH Grant GM030498 to T.M.L. V.M.J was supported by a training grant T32 EB014855 from the National Institute of Biomedical Imaging and Bioengineering, NIH.

## Supplementary Materials

**Fig. S1.**
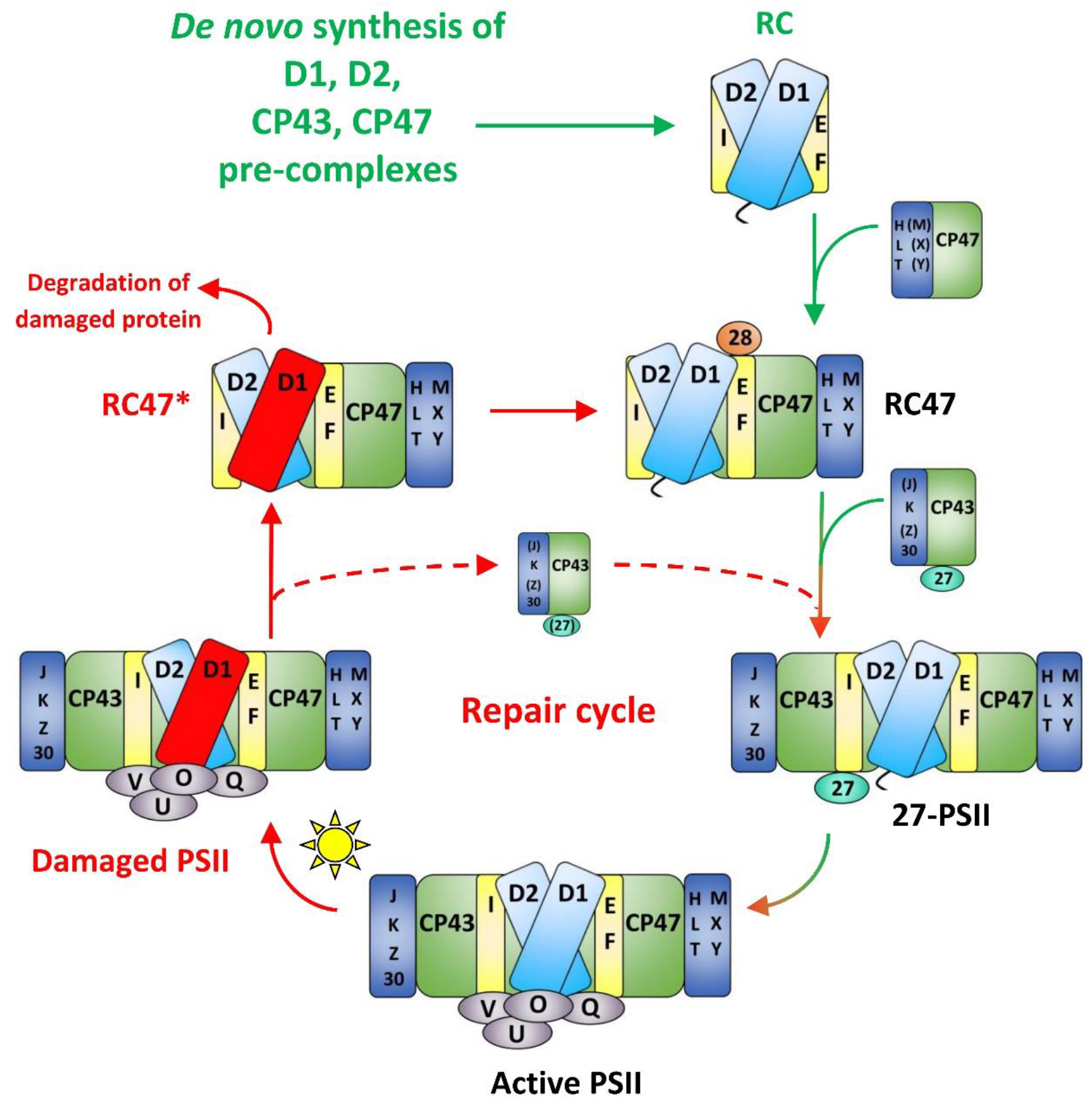
Model of the PSII life cycle based on previous findings [reviewed in (14, 18, 33, 36)]. Several intermediary stages in the life cycle are omitted for clarity; *e.g*., dimerization of active PSII; and after damage, dissociation of PsbO, PsbU, PsbV, and PsbQ (36), Psb27 rebinding (21), and monomerization of the complex (36) before the RC47 formation step. In this model, the damaged RC47 complex serves as the site of D1 replacement. Red D1, damaged D1; RC47*, damaged RC47. Green arrows and text represent *de novo* synthesis steps; red arrows and text represent repair cycle steps. Black text and the red-green arrow represent steps common to both processes.

**Fig. S2.**
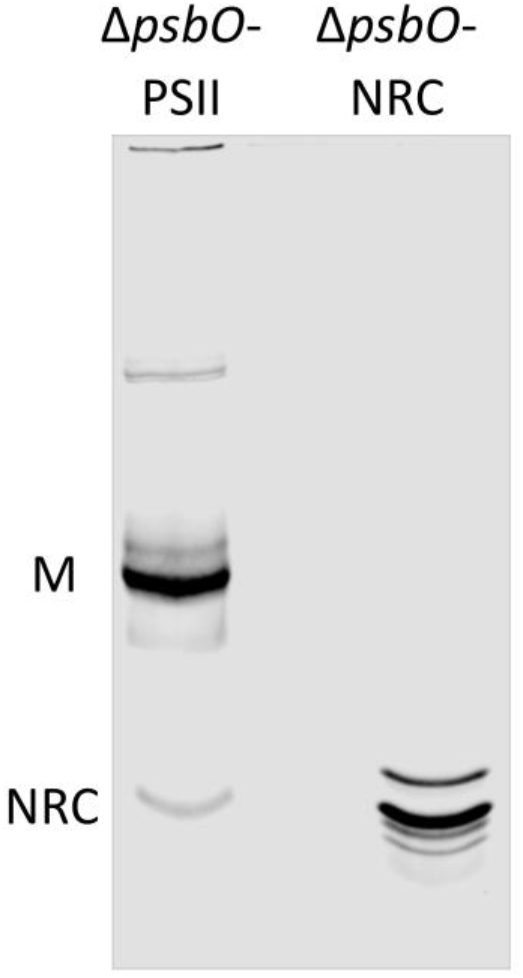
CN PAGE of isolated PSII complexes. Clear Native polyacrylamide gel of *ΔpsbO*-PSII and purified NRC from glycerol gradient of *ΔpsbO*-PSII. The chlorophyll-containing NRC band co-migrates with the unknown band in the PSII sample.

**Fig. S3.**
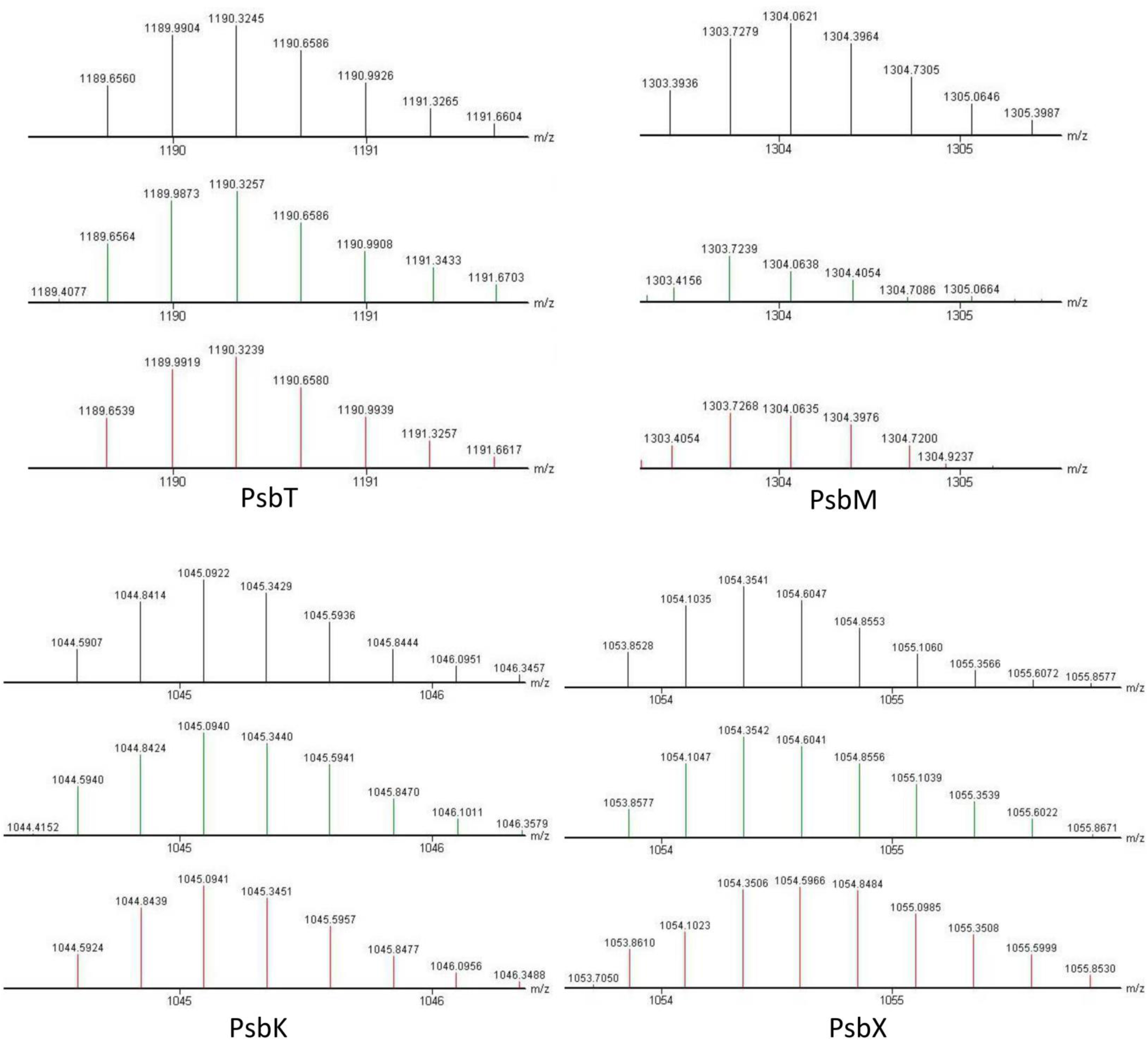
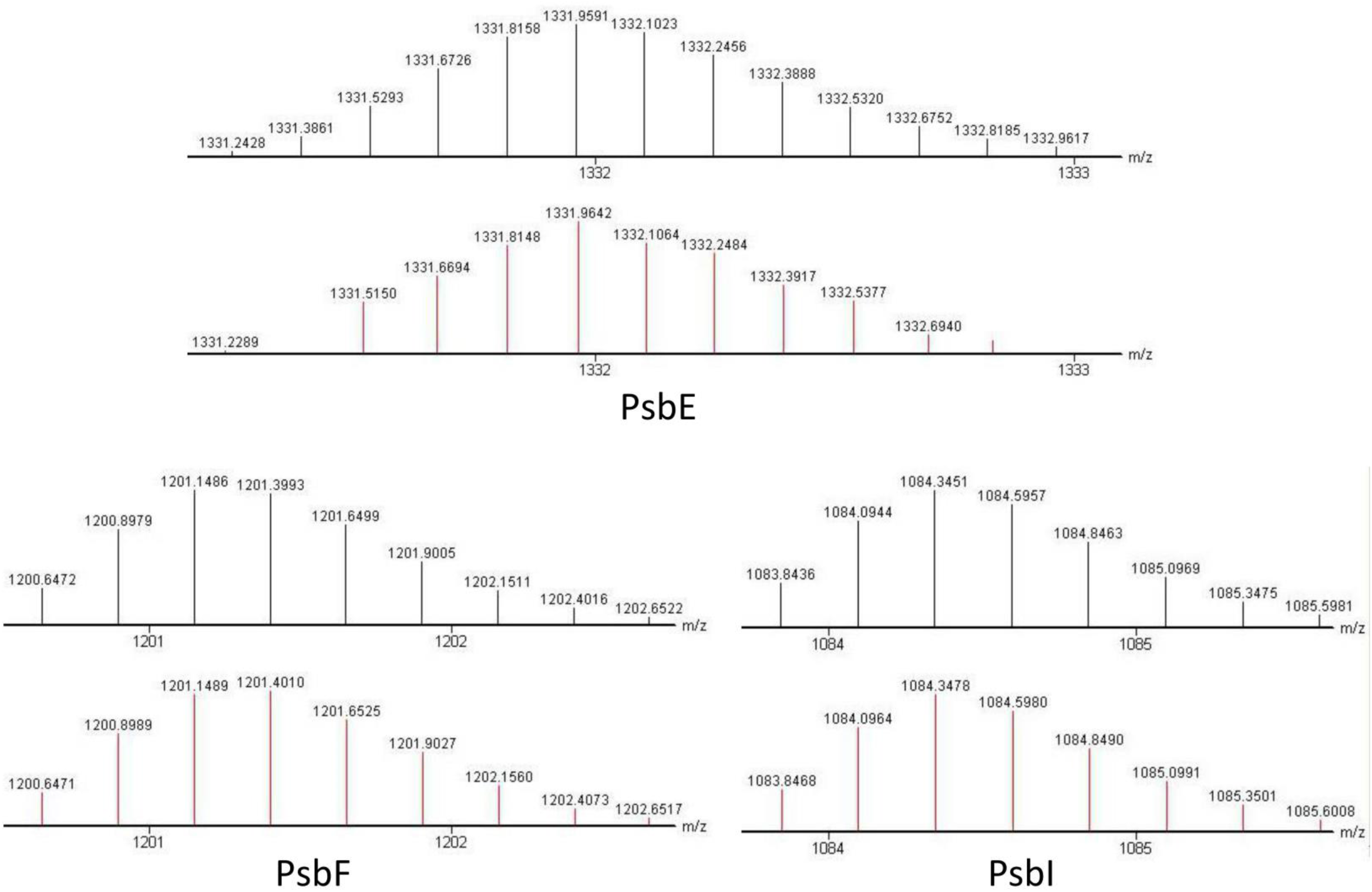
LMM components of PSII complexes. Mass spectra of the intact LMM subunits identified in PSII-M (bottom spectrum for each subunit), NRC (middle), and the theoretical spectrum (top) of each subunit, with protein modifications as listed in Fig. 3.

Tables S1-S5

